# A homozygous missense mutation in *ERAL1,* encoding a mitochondrial rRNA chaperone, causes Perrault syndrome

**DOI:** 10.1101/115790

**Authors:** Iliana A. Chatzispyrou, Marielle Alders, Sergio Guerrero-Castillo, Ruben Zapata Perez, Martin A. Haagmans, Laurent Mouchiroud, Janet Koster, Rob Ofman, Frank Baas, Hans R. Waterham, Johannes N. Spelbrink, Johan Auwerx, Marcel M. Mannens, Riekelt H. Houtkooper, Astrid S. Plomp

## Abstract

Perrault syndrome (PS) is a rare recessive disorder characterized by ovarian dysgenesis and sensorineural deafness. It is clinically and genetically heterogeneous, and previously mutations have been described in different genes, mostly related to mitochondrial proteostasis. We diagnosed three unrelated females with PS and set out to identify the underlying genetic cause using exome sequencing. We excluded mutations in the known PS genes, but identified a single homozygous mutation in the *ERAL1* gene (c.707A>T; p.Asn236Ile). Since ERAL1 protein binds to the mitochondrial 12S rRNA and is involved in the assembly of the small mitochondrial ribosomal subunit, the identified variant represented a likely candidate. *In silico* analysis of a 3D model for ERAL1 suggested that the mutated residue hinders protein-substrate interactions, potentially affecting its function. On a molecular basis, PS skin fibroblasts had reduced ERAL1 protein levels. Complexome profiling of the cells showed an overall decrease in the levels of assembled small ribosomal subunit, indicating that the *ERAL1* variant affects mitochondrial ribosome assembly. Moreover, levels of the 12S rRNA were reduced in the patients, and were fully rescued by lentiviral expression of wild type ERAL1. At the physiological level, mitochondrial respiration was markedly decreased in PS fibroblasts, confirming disturbed mitochondrial function. Finally, knockdown of the *C. elegans ERAL1* homologue *E02H1.2* almost completely blocked egg production in worms, mimicking the compromised fertility in PS-affected women. Our cross-species data in patient cells and worms support the hypothesis that mutations in *ERAL1* can cause PS and are associated with changes in mitochondrial metabolism.

## Introduction

Perrault syndrome (PS) (MIM 233400) is a rare disorder that is inherited in an autosomal recessive manner. It is clinically characterized by sensorineural hearing loss in both male and female patients, while females also present with ovarian dysgenesis, which results in amenorrhea and infertility (1). Some patients also present with neurological manifestations, including ataxia, mild mental retardation and peripheral neuropathy (2–4). Because of the clinical heterogeneity, the disorder has been classified into type I, which is static and without neurological manifestations, and type II that includes progressive neurological symptoms (5). The clinical heterogeneity of PS may partly be due to its genetic heterogeneity; to date, mutations in five different genes have been identified as disease-causing in different cases of PS. The first mutations were reported in *HSD17B4,* a gene encoding a peroxisomal enzyme involved in fatty acid β-oxidation and steroid metabolism (5). Later on, mutations in two genes encoding the mitochondrial aminoacyl-tRNA synthetases HARS2 and LARS2 (6, 7) were identified, which are components of the mitochondrial translation machinery. Finally, two more genes encoding mitochondrial proteins—the peptidase CLPP involved in mitochondrial protein homeostasis (8, 9) and the helicase Twinkle that is required for mitochondrial DNA (mtDNA) maintenance (10)—were found mutated in some PS patients. Altogether, four out of five PS causing genes are involved in mitochondrial function, particularly protein homeostasis, suggesting that mitochondria play a critical role in the development of the disease.

Here, we report a homozygous c.707A>T (p.Asn236Ile) missense mutation in the *ERAL1* gene (NM_005702.2) identified by exome sequencing in two unrelated patients, and found a third PS patient with the same variant during the course of our study. The human ERAL1 protein (UniProtKB O75616) has been described as an rRNA chaperone indispensable for the assembly of the small 28S subunit of the mitochondrial ribosome (11, 12). In line with this, patient skin fibroblasts and *C. elegans* with knockdown of the *ERAL1* homologue display mitochondrial dysfunction, strongly suggesting that the identified mutation in *ERAL1* is the cause of PS in our patients.

## Results

### Identification of a homozygous missense mutation in the ERAL1 gene in three PS patients

Two women of Dutch ancestry, unrelated but both from the same village, presented at our clinic with symptoms of PS. In PS patient 1 (aged 66 years) hearing loss was diagnosed when she was 20 years old, but probably started much earlier and appeared to be progressive. She had normal menarche at 11 years of age, with irregular menses until the age of 27, when menopause occurred. One sister also had sensorineural hearing loss and premature ovarian failure, but did not participate in our study. PS patient 2 (aged 38 years) was diagnosed at 4 years of age with sensorineural hearing loss, which was more severe in the high frequencies and slowly progressive. At the age of 18 years she presented with primary amenorrhea and underdeveloped secondary sexual characteristics. Abdominal ultrasound revealed streak ovaries and a small uterus. An ovary biopsy showed fibrous tissue without primordial follicles. Her father had sensorineural hearing loss since childhood, but no fertility problems. Her mother and two sisters were healthy.

In order to identify the underlying genetic cause for the disease, we performed whole exome sequencing (WES), which excluded the presence of mutations in the known PS genes *HSD17B4* (5), *HARS2* (6), *LARS2* (7), *CLPP* (8) and *C10orf2* (10). Because both patients originate from a small village, a known genetic isolate, recessive inheritance and shared genetic cause were suspected. Subsequent filtering of all variants found in WES for homozygous variants that were shared in both patients, and with a minor allele frequency of <1% in public or in house databases yielded only one variant: a homozygous nucleotide substitution at position c.707A>T (p.Asn236Ile) in *ERAL1* (NM_005702.2), a gene encoding the Era-like 12S mitochondrial rRNA chaperone 1.

The substituted residue is highly conserved in vertebrates as well as in fruit flies and in *C. elegans* (Fig. 1A). Moreover, the variant is predicted to be “probably damaging” by PolyPhen-2 (HumDiv score 1.000, HumVar score 0.992), SIFT (score 0) and MutationTaster (p-value 1). Subsequent sequencing of family members of patient 2 showed that the healthy females are heterozygous while the affected father is homozygous for this variant (Fig. 1B). The identified variant is not present in any public database. However, from the genotyping of 530 individuals of the same village as the patients, 49 were found heterozygous and 0 homozygous for the variant, demonstrating an allele frequency of 4.6% in this genetic isolate. This would suggest a relatively high prevalence (0.2%) of PS in this village. Indeed, during the course of our study a third PS patient, presenting with progressive sensorineural hearing loss since birth and primary amenorrhoea, visited our outpatient clinic and was found to be homozygous for the same *ERAL1* variant (Fig. 1C).

**Figure 1.**
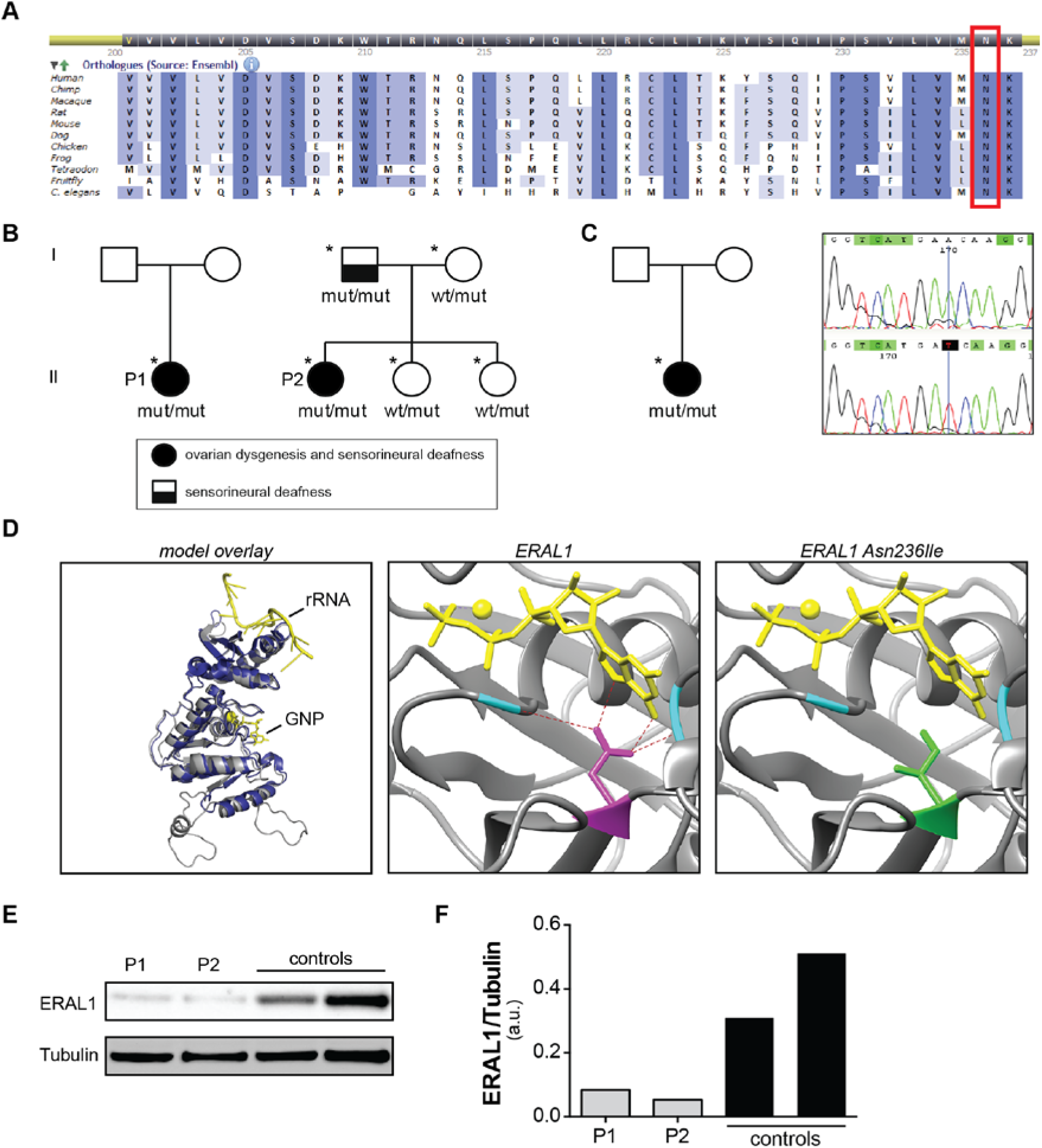
Three individuals diagnosed with PS, carry the same homozygous missense mutation in the *ERAL1* gene. (A) The mutated residue is highly conserved as evidenced by sequence alignment using the Alamut software. (B) Pedigrees of the two initial PS patients (indicated by P1 and P2, as annotated in the remainder of the paper) in whom WES was performed and skin fibroblasts were analyzed. Individuals that were sequenced for the mutation are indicated with asterisks, and the results are annotated (wt: wildtype ERAL1, mut: mutated ERAL1). (C) Pedigree (left panel) and sequencing results (right panel) of a third PS patient that was found homozygous for the same mutation. (D) Left panel: Structural Alignment of modelled human ERAL1 (gray) and crystallized Aquifex aeolicus ERA (dark blue), showing the high similarity between the two structures. Ligands of ERA are shown in yellow. Middle panel: Active center of modelled wild type ERAL1 in complex with a non-hydrolizable GTP analog (GNP, phosphoaminophosphonic acid-guanylate ester) (yellow sticks) and magnesium (yellow sphere). Hydrogen bonds between asparagine 236 (magenta sticks), GNP and alanines 124 and 310 (cyan) are shown as red dashes. Right panel: Active center of modelled N236I mutated ERAL1 in complex with GNP (yellow sticks) and magnesium (yellow sphere), showing that no interactions can be made between the mutated amino acid, the ligand and the two flanking alanines (cyan). (E) Western blot of skin fibroblasts from PS patients and controls; both PS patients present with decreased ERAL1 protein levels. P1: PS patient 1, P2: PS patient 2. (F) Bar graph depicting the levels of ERAL1 in patient and control fibroblasts normalized to tubulin, as quantified from the blot in panel E.

ERAL1 protein is indispensable for the assembly of the small 28S subunit of the mitochondrial ribosome through binding to its 12S rRNA component (11, 12), and is therefore involved in the translation of mtDNA-encoded proteins. To predict *in silico* whether the p.Asn236Ile variant could affect the rRNA-ERAL1 interaction, we constructed a 3D model of the protein based on the crystal structure of the bacterial orthologue ERA, which has been characterized in the bacterium *Aquifex aeolicus* (13). We performed a 3D structural alignment between the bacterial ERA crystal and the human ERAL1 model in order to visualize the position of the residue found mutated in our patients in comparison to its bacterial counterpart (Fig. 1D, left panel). The alignment showed that the position of the asparagine 236 (N236) in human ERAL1 corresponds to the *Aquifex aeolicus* ERA asparagine 123 (N123), which has been described to directly interact with GTP through the formation of hydrogen bonds (13) (Fig. 1D, middle panel). Additionally, ERA N123 interacts with two amino acids, valine 14 (V14) and alanine 156 (A156) (corresponding to the ERAL1 residues A124 and A310), that are part of the loops flanking the substrate and, therefore, crucial for maintaining the active center’s integrity (Fig. 1D, middle panel). Binding of ERA with GTP is necessary for its function, causing changes in its conformation, followed by further interaction of ERA-GTP with the bacterial 16S rRNA (13). Substitution of the polar amino acid asparagine with the hydrophobic isoleucine at position 236 in the human ERAL1 (Fig. 1D, right panel) is likely to impair interactions with GTP and the A124 and A310 residues, and predicted to disrupt protein conformation and its interaction with the human 12S rRNA.

Because previously described mutations causing PS are mostly in genes related to mitochondrial homeostasis—two of which are directly involved in mitochondrial translation (6, 7), one in mtDNA maintenance (10) and one in mitochondrial proteostasis (8)—we hypothesized that the identified *ERAL1* variant is likely to cause PS in our patients. To test this hypothesis we set out to investigate the effects of the ERAL1 variant at a cellular as well as at an organismal level.

### ERAL1 protein levels and assembly of the 28S ribosomal subunit are compromised in PS patients

To test whether the sequence variant identified in our patients affects ERAL1 protein levels, we performed a Western blot on lysates from cultured skin fibroblasts of PS patients and control subjects. We observed that in both PS patients, ERAL1 protein levels where decreased when compared to fibroblasts from healthy controls (Fig. 1E and F).

Since ERAL1 is involved in the assembly of the small 28S mitochondrial ribosomal subunit (11, 12), we next investigated whether the PS patient cells showed impaired assembly of the 28S subunit. We assessed the abundance of assembled mitochondrial ribosomal subunits using complexome profiling (14). With this technique migration profiles and relative abundance of protein complexes are identified using a combination of blue native electrophoresis and shotgun proteomics. Although the migration profiles of the proteins composing the small (28S) and large (39S) mitochondrial ribosomal subunits were similar between PS patients and controls (Fig. 2A and B), we observed a remarkable decrease (30-40%) in the overall abundance of proteins composing the small 28S subunit in the PS patients compared to healthy controls (Fig. 2A and C). In contrast, the abundance of proteins composing the large 39S subunit was comparable between cells from PS patients and controls (Fig. 2B and D). These observations led us to conclude that the overall abundance of assembled small mitochondrial ribosomal subunits is lower in cells from the PS patients compared to those from healthy controls, suggesting that the identified ERAL1 variant perturbs proper assembly of the small 28S mitochondrial ribosomal subunit.

**Figure 2.**
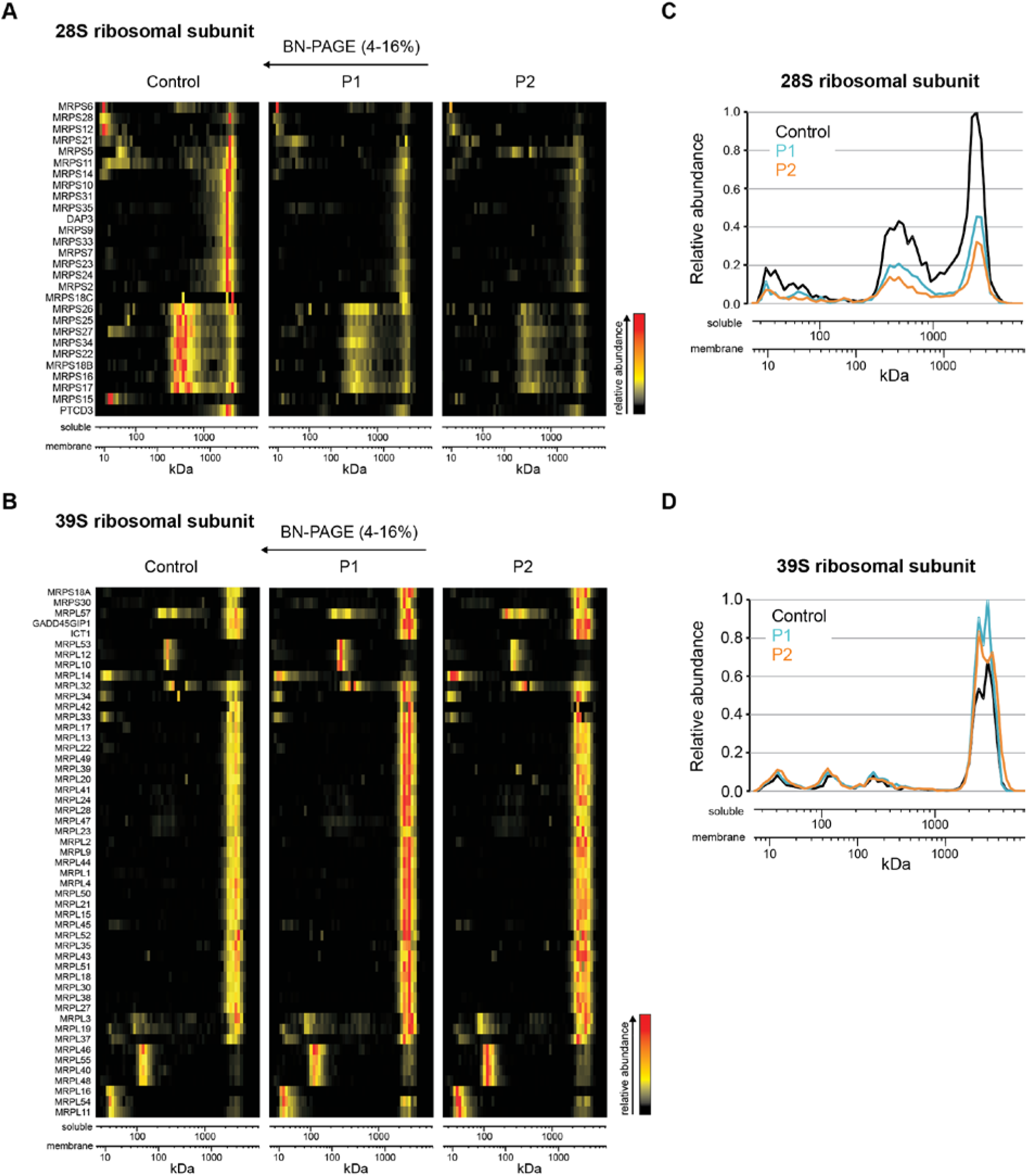
Cells from PS patients show defected assembly of the small 28S mitochondrial ribosomal subunit. (A-B) Heat map representation of migration profiles in blue native gels of proteins of the small 28S (A) and large 39S (B) mitochondrial ribosomal subunits isolated from PS patient and control skin fibroblasts. (C-D) Graphs depicting the average normalized relative abundance of proteins of the 28S (C) and 39S (D) mitochondrial ribosomal subunits spanning the blue native gel. Protein abundance was determined by label-free quantitation using the composite iBAQ intensity values determined by MaxQuant (31) and normalized considering multiple migration profiles of individual proteins, that is taking into account iBAQ values from all 180 gel slices (60 slices per sample). Both patients show decreased levels of assembled small mitochondrial ribosomal subunit (A,C), while the levels of assembled large subunit remain unaffected (B,D). P1: patient 1, P2: patient 2.

### 12S rRNA levels are low in PS patients, and rescued to control levels after ERAL1 lentiviral expression

Because ERAL1 acts as a chaperone to the mitochondrial 12S rRNA of the small ribosomal subunit by protecting it from degradation (12), we hypothesized that levels of 12S rRNA are reduced in the patients, while the 16S rRNA of the large ribosomal subunit remains unaffected. We therefore employed qPCR and measured the 12S/16S rRNA ratio in patient and control fibroblasts. In support of our hypothesis we found that this ratio was significantly reduced in the two patients compared to healthy controls (Fig. 3A).

Next, we asked whether ectopic expression of wild type ERAL1 in the patient cells would be sufficient to reconstitute 12S rRNA levels. To address this question, we infected patient fibroblasts with lentiviral particles overexpressing either ERAL1 or GFP as a control. We found that overexpression of ERAL1 in both patients (Fig. 3B) fully rescued the 12S/16S rRNA (Fig. 3C). These data demonstrate that the identified mutation leads to reduced 12S rRNA levels, a condition that is reversible after overexpression of wild type ERAL1.

**Figure 3.**
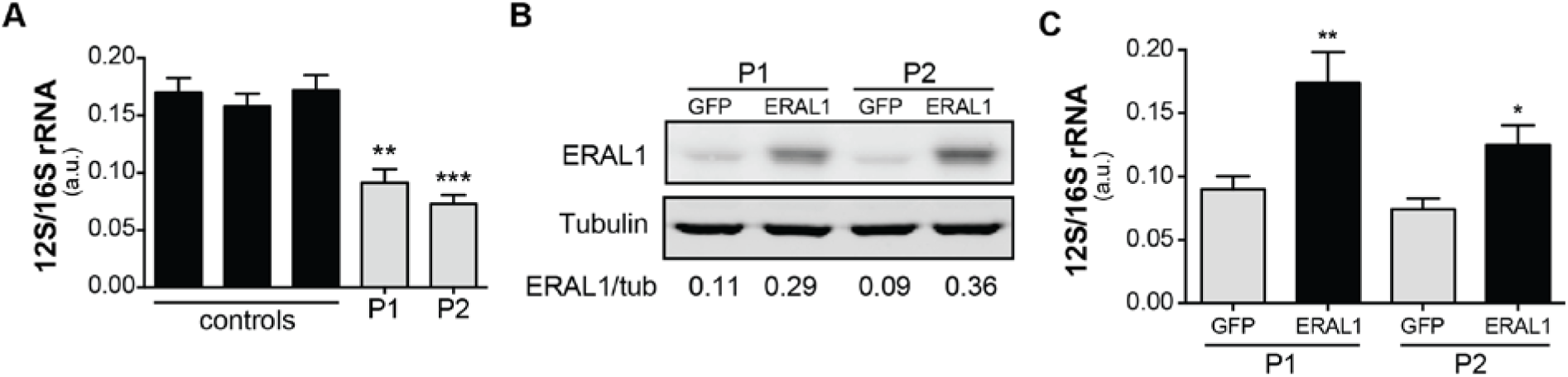
12S rRNA levels are decreased in PS cells and are fully rescued after ectopic ERAL1 expression. (A) The 12S to 16S rRNA ratio in control and PS fibroblasts as measured by qPCR. Both patients present with significantly lower 12S/16S rRNA ratio compared to controls. Data are represented as the mean±SEM of six biological replicates. **p<0.01, ***p<0.001 as calculated by a one-way ANOVA Tukey's multiple comparisons test. (B) Western blot from PS fibroblasts infected by GFP or ERAL1 overexpressing lentivirus. Patient cells infected with ERAL1 overexpressing lentivirus stably express ERAL1 protein compared to those infected by a GFP overexpressing virus. Quantification of the levels of ERAL1 normalized to tubulin are given below the blot. (C) 12S to 16S rRNA ratio in PS fibroblasts stably expressing ERAL1 or GFP. Patient cells stably expressing ERAL1 present a significantly increased 12S/16S rRNA ratio compared to those expressing GFP, and reaches the levels of healthy cells (panel A). Data are represented as the mean±SEM of six biological replicates. *p<0.05, **p<0.01 as calculated by a 2-tailed student’s t-test.

### Mitochondrial function is impaired in PS patients

Disturbed assembly of the mitochondrial ribosome may lead to inefficient translation of mitochondrial DNA (mtDNA)-encoded proteins, which form core subunits of the oxidative phosphorylation (OXPHOS) complexes. To test whether mitochondrial translation is disturbed, we quantified the ratio between the mtDNA-encoded cytochrome *c* oxidase subunit 1 (MT-CO1) and the nuclear DNA (nDNA)-encoded succinate dehydrogenase complex, subunit A (SDHA) (15) in the PS cells. Indeed, the abundance of MT-CO1 was found lower, while SDHA was unaffected (Fig. 4A and B). We next measured mitochondrial activity by means of oxygen consumption rate (OCR) in the cells using Seahorse respirometry. Basal respiration was decreased in the PS cells when compared to controls (Fig. 4C). After injection of the uncoupler FCCP, maximal respiratory capacity of the PS fibroblasts was also impaired (Fig. 4C). These findings confirm that mitochondrial function is compromised in the PS cells.

**Figure 4.**
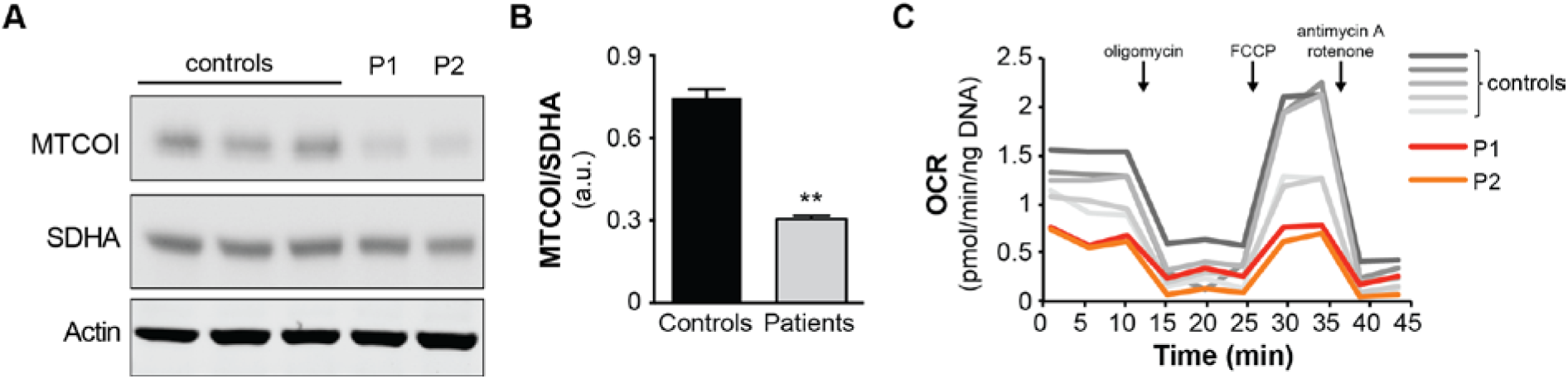
Cells from PS patients show impaired mitochondrial function. (A) Western blot from PS patient and control skin fibroblasts; both patients present with decreased MT-CO1 protein levels, while SDHA levels are unaffected. P1: PS patient 1, P2: PS patient 2. (B)Histogram depicting the MT-CO1 to SDHA ratio, as quantified from a western blot with samples depicted in panel A, in biological duplicates. Data are represented as the mean±SEM. **p<0.01 as calculated by a 2-tailed student’s t-test. (C) Seahorse respirometry on PS patient and control skin fibroblasts; both PS patients have reduced basal and maximal respiration. Injections of compounds during measurement are indicated with arrows. OCR: oxygen consumption rate, P1: PS patient 1, P2: PS patient 2.

### Knockdown of the ERAL1 homologue in C. elegans compromises fertility

In order to test the phenotypic effects of low ERAL1 levels at an organismal level, we used the nematode *C. elegans* as a model organism and specifically knocked down the expression of the worm *ERAL1* homologue *E02H1.2.* We used the RNAi-sensitive nematode strain *rrf-3(pk1426)* (16). Worms exposed to *E02H1.2* RNAi had approx. 50% knockdown (Fig. 5A), developed normally and had normal lifespan compared to control HT115 worms (data not shown). Strikingly, we observed that the animals undergoing *E02H1.2* RNAi were not carrying eggs throughout adulthood (Fig. 5B). We quantified the number of eggs laid during the first four days of adulthood, which is the fertile period of *C. elegans,* and confirmed that the *E02H1.2* RNAi fed animals hardly laid any eggs (Fig. 5C).

**Figure 5.**
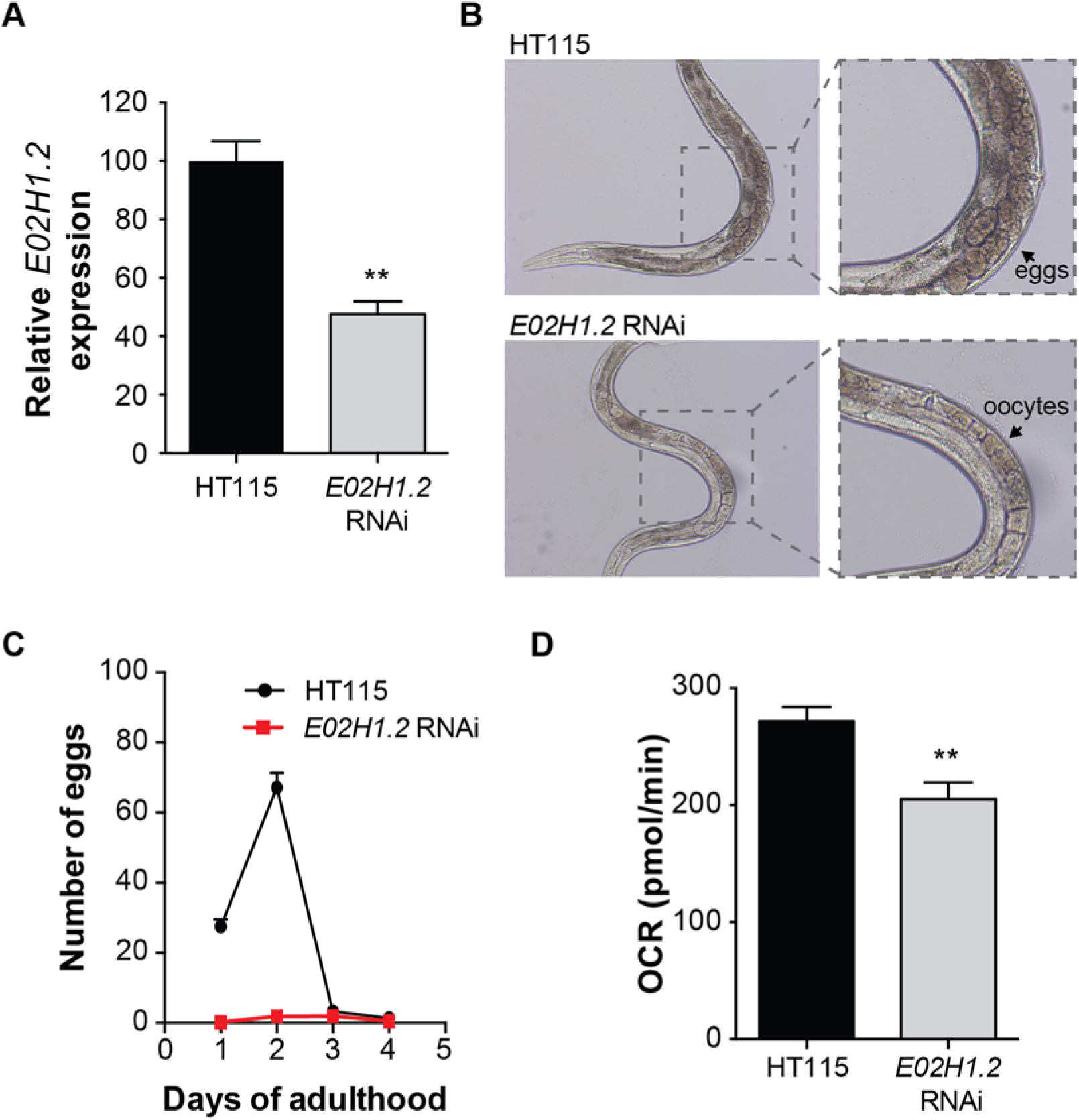
Knockdown of the ERAL1 worm homologue E02H1.2 compromises fecundity and reduces respiration in C. elegans. (A) mRNA levels of knockdown efficiency of E02H1.2 in worms fed with control HT115 and E02H1.2 RNAi as measured by qPCR. Worms fed with E02H1.2 RNAi present with 50% reduction of E02H1.2 gene expression. Data are represented as the mean±SEM of biological triplicates. **p<0.01 as calculated by a 2-tailed student’s t-test. (B) Wide-field images of adult (day 1) worms fed with control HT115 (upper panels) and E02H1.2 (lower panels) RNAi; worms with E02H1.2 knockdown do not carry any eggs, in contrast to the controls. (C) Graph showing the number of eggs laid over the first four days of adulthood. Worms with E02H1.2 knockdown laid almost no eggs compared to the controls. Egg-laying of 30 individual worms was followed. Error bars correspond to SEM. (D) Basal respiration measured on worms fed with control (HT115) or E02H1.2 (ERAL1) RNAi. Worms with E02H1.2 knockdown have significantly (p<0.01) decreased basal respiration compared to the controls. n=8. Error bars correspond to SEM.

Because the PS skin fibroblasts presented with decreased mitochondrial respiration, we also measured OCR in the worms with *E02H1.2* knockdown. Those worms demonstrated significantly decreased OCR (Fig. 5D). Collectively, these results demonstrate that low ERAL1 expression impairs mitochondrial function and compromises fecundity in *C. elegans,* mimicking the defects we observe in our PS patients.

## Discussion

Three women from three different families presented with signs of PS, including deafness and ovarian dysgenesis. We employed exome sequencing and identified a homozygous missense mutation in the *ERAL1* gene, c.707A>T. The mutation leads to a damaging p.Asn236Ile substitution that is likely to interfere with the GTP binding capacity of the ERAL1 protein. Specifically, the observation that the assembly of the small mitochondrial ribosomal subunit—for which ERAL1 is known to play a pivotal role (11, 12)—is compromised in the PS cells, implies that the identified mutation affects proper function of ERAL1 protein. Importantly, the impaired 12S rRNA levels in PS fibroblasts and their full rescue upon expression of wild type ERAL1 underlines the causal link between the mutation and mitochondrial function. Additionally, the decreased respiration that was noted in the PS fibroblasts demonstrates disturbed mitochondrial function, confirming that the identified mutation affects mitochondria and is pathogenic. Finally, we performed RNAi experiments in the nematode *C. elegans* to demonstrate the role of ERAL1 in fertility. Knockdown of the worm *ERAL1* homologue *E02H1.2* indeed impaired egg production presenting a crucial role of this protein in the nematode’s fecundity. Our cross-species findings on both patient fibroblasts and the model organism *C. elegans* identified the *ERAL1* variant as the cause for the clinical symptoms in our three PS patients.

The identification of *ERAL1* as a causative gene for PS extends the mutational spectrum for this disease. So far, mutations in five different genes have been described to cause Perrault syndrome. These include the *HARS2* (6) and *LARS2* (7) genes, both encoding mitochondrial tRNA synthetases, the *CLPP* (8) that codes for a protease of the mitochondrial matrix and the *C10orf2* (10) encoding the mitochondrial helicase Twinkle. These findings imply that mitochondria are involved in the development of the disease. Interestingly, recent work on *CLPP* deficient mice (17) elegantly links ERAL1 mechanistically to CLPP by demonstrating that removal of ERAL1 from the assembled 28S ribosomal subunit is essential for the final maturation of the whole 55S mitochondrial ribosome, and that this removal requires CLPP. Our identification of ERAL1 as a Perrault syndrome gene further ties CLPP and ERAL1 together, as mutations in either of them lead to similar pathology in humans.

The involvement of mitochondrial dysfunction in development of deafness is already well established (18–20). On the other hand, the role of mitochondria in infertility is less known and is only recently starting to emerge. Studies in mice have shown that perturbations of mitochondrial function can affect oocyte maturation (21, 22). In humans, age-related decline in oocyte quality and quantity has been associated with mitochondrial dysfunction and impaired OXPHOS, while dietary supplementation with the OXPHOS enhancer CoQ10 improved ovulation in aged women (23). In *C. elegans,* knockdown of the mitochondrial tRNA synthetase *hars-1* led to smaller gonads and to loss of fertility, which was partly due to germ cell apoptosis (6). Similarly, the *lars-2* mutant worm had underdeveloped gonads and was completely sterile (7). Also, sub-lethal disruptions in OXPHOS subunits cause sterility in worms (24–26). Our findings of compromised fertility and mitochondrial respiration after knockdown of the worm *ERAL1* homologue serve as additional proof for the importance of mitochondria in fertility. The exact mechanisms through which mitochondria regulate fertility, however, remain to be resolved.

Collectively, our results indicate that mutations in another mitochondrial gene, *ERAL1,* can cause sensorineural deafness and ovarian dysgenesis of PS, strengthening the notion that this syndrome is caused by dysfunctional mitochondria. Detailed functional mitochondrial assays with cells from other unresolved cases of PS may shed more light on the molecular mechanisms underlying the disease and reveal other PS-causing genes.

## Material and Methods

All three PS patients visited the outpatient clinic of the AMC and signed informed written consent for this study. Information regarding the medical history was obtained by interviewing the patients and from their medical files.

### Whole Exome Sequencing (WES)

WES of PS patients was conducted using the SeqCap EZ Human Exome Library v3.0 (Roche NimbleGen) and a 5500 SOLiD™ instrument (Life Technologies). Samples were prepared using standard SolID 75×35 paired end sequencing protocols. Alignment of sequence reads to human reference genome (hg19) was done using Lifescope 2.5.1, and variants were called using the GATK2.5 software package. Mean target coverage was 86x for PS patient 1 and 66x for PS patient 2. Coverage of targeted regions at 10x read depth (after removal of duplicate reads) was 87% for patient 1 and 84% for patient 2.

Prioritization of variants identified with WES was done using the Cartagenia BENCHlab NGS software (Cartagenia NV). Public databases used for determining the frequency of the identified variants in the general population were: 1000 genomes (1000 Genomes Phase 3 release v5.20130502), dbSNP (dbSNP build 141 GRCh37.p13), the ESP6500 dataset (http://evs.gs.washington.edu/EVS/), and the GoNL database (498 Dutch individuals, http://www.nlgenome.nl/). Variants were further characterized using Alamut version 2.3 (Interactive Biosoftware, Rouen, France).

### Sanger sequencing of ERAL1

Confirmation of the mutation in the PS patients and genotyping of family members of PS patient 2 was done by Sanger sequencing. Primers were designed to amplify exon 6 of *ERAL1* using the Primer3 software (http://bioinfo.ut.ee/primer3-0.4.0/). Amplification was performed with M13-tagged primers using HOT FIREPol™ DNA polymerase (Solys Biodyne) and a touchdown PCR program. PCR fragments were sequenced using the Bigdye kit v1.1 (Applied Biosystems). Reactions were run on an ABI3700 or ABI3730XL genetic analyzer (Applied Biosystems) and sequences were analyzed using Sequence Pilot (JSI Medical systems) or CodonCode aligner (CodonCode Corporation).

### Genotyping TaqMan

Genotyping of 530 individuals from the same village as the PS patients was done by a TaqMan assay on a Roche LC480 lightcycler. For the SNPs, Sanger sequencing confirmed homozygous reference, heterozygous and homozygous mutant samples were used as standards for SNP calling.

### ERAL1 3D model

The crystallized structure of *Aquifex aeolicus* ERA (PDB code: 3IEV) was selected as a template to build a model for human ERAL1 using SWISS-MODEL server (http://swissmodel.expasy.org/) (27), as it is the most similar crystallized structure available in the database (32% identity, 0.35 similarity). UCSF Chimera package (Version 1.10.2, Computer Graphics Laboratory, University of California, San Francisco) was used to align and compare the two 3D structures in order to predict the possible implications of the mutated amino acid on the function of ERAL1 protein.

### Cell culture

Primary skin fibroblasts from PS patients and healthy controls were cultured in DMEM or Ham’s F-10 with L-glutamine (Bio-Whittaker) growth media, supplemented with 10% fetal bovine serum (Bio-Whittaker), 25 mM HEPES buffer (BioWhittaker), 100 U/ml penicillin, 100 μg/ml streptomycin (Life Technologies), and 250 ng/ml Fungizone (Life Technologies) in a humidified atmosphere with 5% CO_2_ at 37°C.

### Immunoblot analysis

For protein extraction, cultured skin fibroblasts were lysed in RIPA buffer (50 mM Tris-HCl pH7.4, 150 mM NaCl, 0.1% sodium dodecyl sulfate, 0.5% Sodium deoxycholate, 1% Triton X-100) with addition of Complete mini protease inhibitor cocktail (Roche). Samples were sonicated to ensure complete lysis, and briefly centrifuged to discard debris. Protein concentrations were measured using the BCA protein assay kit (Pierce). For immunoblot analysis, lysates were diluted in NuPAGE LDS Sample Buffer and Sample Reducing Agent (Life Technologies) and heated to 70°C. Protein extracts were separated on pre-cast NuPAGE 4-12% Bis-Tris gels (Life Technologies), and transferred to a nitrocellulose membrane. Membranes were blocked with 3% (w/v) milk powder in PBS containing 0.1% (v/v) Tween-20 (PBS-T), and incubated overnight at 4°C with the primary antibody. The following day membranes were washed with PBS-T and incubated for 1 h at room temperature with the secondary antibody. Odyssey imaging system (LI-COR) was used for imaging of membranes incubated with IRDye secondary antibodies (LI-COR). Membranes incubated with HRP-linked secondary antibodies were detected using the ECL prime western blotting detection reagent (Amersham) and imaged with the ImageQuant LAS4000 (GE Healthcare). Primary antibodies: ERAL1 (Proteintech 11478-1-AP), MTCO1 (Abcam ab14705), SDHA (Abcam ab14715), Tubulin (Sigma T6199), Actin (Abcam ab14128). Secondary antibodies: Clean-Blot IP Detection Reagent HRP secondary antibody (Thermo Scientific 21230), Goat anti-mouse-HRP (Dako P0446), IRDye 800CW Goat-anti-Mouse (LI-COR 926-32210), IRDye 680RD Donkey-anti-Mouse (LI-COR 926-68072). Quantification of bands was performed using the AIDA software.

### Isolation of mitochondria

Isolation of mitochondria was carried out as previously described (28) with minor modifications. Cell pellets were resuspended in 125 mM sucrose, 1 mM EDTA, 20 mM Tris/HCl pH 7.4 and disrupted by 10 strokes in a glass/Teflon Potter-Elvehjem homogenizer.

The homogenates were mixed with 875 mM sucrose, 1 mM EDTA, 20 mM Tris/HCL pH 7.4 to adjust the sucrose concentration and centrifuged at 1000 *g* for 10 min at 4 °C. Supernatants were centrifuged at 14000 *g* for 10 min at 4°C. The mitochondrial pellets were resuspended in 250 mM sucrose, 1 mM EDTA, 20 mM Tris/HCl pH 7.4. Protein concentration was determined by the Lowry method.

### Quantitative PCR (qPCR)

For qPCR, RNA was isolated using the TRIzol reagent (Invitrogen) and cDNA synthesis was performed with 1μg RNA using the QuantiTect Reverse Transcription Kit (QIAGEN). qPCR was performed on a Roche Lightcycler 480 using Roche SYBR-green mastermix. For measurement of the mitochondrial ribosomal RNAs, cDNA samples were heated for 10 min at 95 °C, followed by 36 cycles of 15 sec at 95 °C (denaturation), 10 sec at 56 °C (annealing) and 15 sec at 72 °C (amplification). To ensure that no genomic rRNA is amplified, samples without reverse transcriptase were also included in the qPCR run. For measurement of *E02H1.2* knockdown efficiency in *C. elegans,* cDNA samples were heated for 6 min at 95 °C, followed by 40 cycles of 10 sec at 95 °C (denaturation), 5 sec at 65 °C (annealing) and 15 sec at 72 °C (amplification). *E02H1.2* transcript levels were normalized to the geometrical mean of three different house keeping genes *(CDC42, ACT, F35G12.2).*

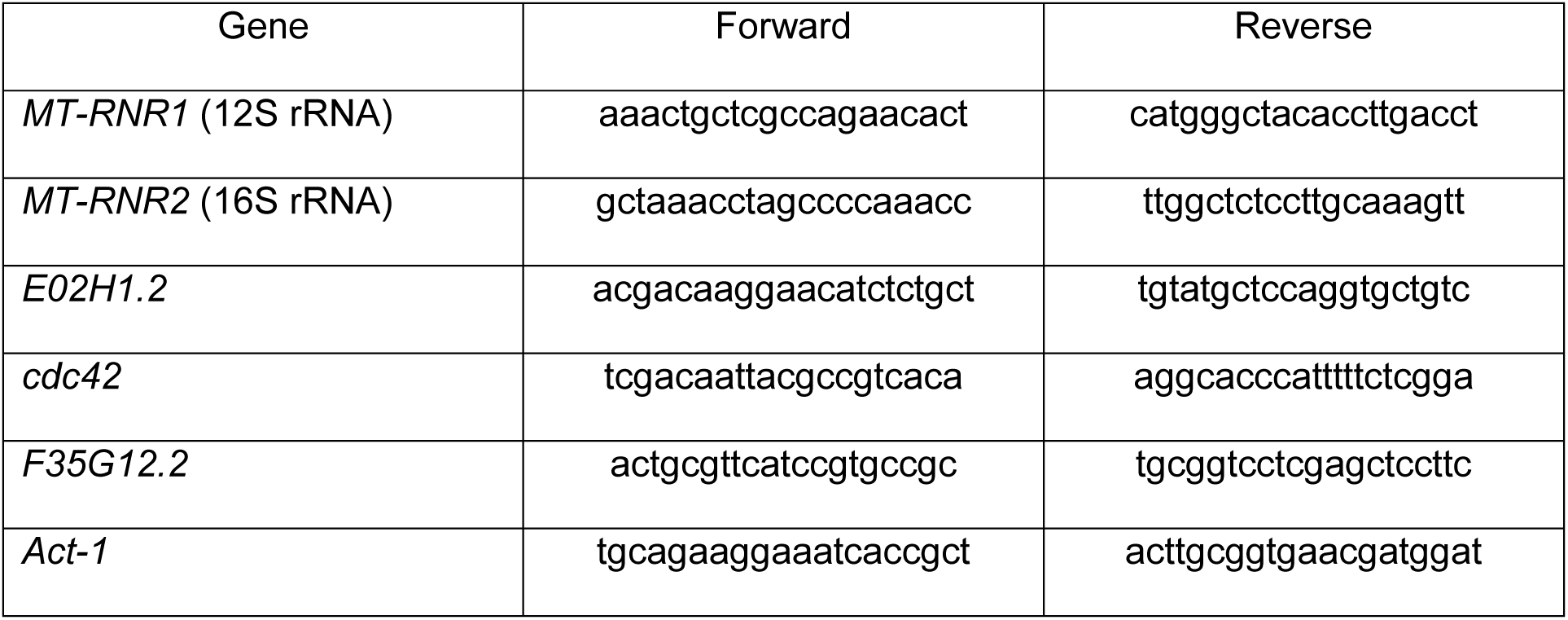
List of primers:

### Cloning and viral transfection

For the functional complementation assay, cells were infected with viral particles for stable overexpression of either ERAL1 or GFP (pLenti/ERAL1, pLenti/GFP). The pLenti/ERAL1 vector was constructed using the Gateway technology (Invitrogen). Specifically, the ERAL1 ORF PCR product containing attB sites was cloned to the entry plasmid pDONR 221 (Invitrogen) using the BP Clonase enzyme mix (Invitrogen) and further cloned to the destination vector pLENTI 6.3/TO/V5-DEST (Invitrogen) using the LR Clonase enzyme mix (Invitrogen). The pLenti/GFP vector was a kind gift of Prof. Dr. Noam Zelcer (Department of Biochemistry, Academic Medical Center, University of Amsterdam). For virus production, HEK293 cells at ~50% confluency were transfected with the ERAL1 or GFP overexpressing plasmids, together with the lentiviral packaging plasmids pMD2G, pMDL/RRE, pRSV/REV using the DNA and siRNA transfection reagent jetPRIME^®^ (VWR) in normal DMEM culture medium. The following day the medium was refreshed, and viral supernatant was collected and filtered 48 and 72 hours post transfection. Patient fibroblasts at ~70% confluency were infected with the viral supernatant, grown under blasticidine selection (10μg/mL) for 2 weeks, and further expanded in blasticidine-free medium for 4 more passages. Cells were regularly checked for GFP expression. Overexpression of ERAL1 was confirmed by Western blot analysis.

### Complexome profiling

Mitochondrial pellets (200 μg protein) were solubilized with 6 mg digitonin/mg protein and separated by 4-16% gradient Blue native PAGE (BN-PAGE) (29). Complexome profiling was done according to Heide *et al.*(14). Protein identification and data analysis was done essentially as previously described (30) using MaxQuant (version 1.5.0.25 (31)).

Protein abundancies were determined by label-free quantitation using the composite iBAQ values determined by MaxQuant (31) and normalized considering multiple migration profiles of individual proteins, that is taking into account iBAQ values from all 180 gel slices (60 slices per sample). The profiles were hierarchically clustered by distance measures based on Pearson correlation coefficient (uncentered) and the average linkage method and further analyzed by manual correlation profiling. The clustering and the visualization and analysis of the heat maps were done with the NOVA software v0.5 (32).

### Respiration assays

Oxygen consumption was measured using the Seahorse XF96 analyzer (Seahorse Bioscience). Primary skin fibroblasts were plated in 96-well Seahorse plates at a density of 10,000 cells per well and incubated overnight under normal cell culture conditions. The following day, medium was replaced by DMEM (Sigma, D5030) containing 25 mM glucose (Sigma), 1 mM sodium pyruvate (Lonza), and 2 mM L-Glutamine (Life technologies). Basal respiration was measured 3 times, followed by measurements after addition of 1.5 μM oligomycin, 1 μM FCCP and 2.5 μM antimycin A and 1.25 μM rotenone. For respiration assays in worms, animals were transferred in 96-well Seahorse plates (20 worms per well) and basal oxygen consumption was measured 6 times.

### C. elegans culturing and RNAi experiments

The *C. elegans* RNAi-sensitive strain *rrf-3(pk1426)* was obtained from The Caenorhabditis Genetics Center (CGC, University of Minnesota). Worms were cultured and maintained as described previously (33) in 20°C incubators and fed with *E. coli* OP50 strain, also obtained from the CGC. RNAi experiments were carried out as previously described (34). The RNAi clone targeting the ERAL1 worm orthologue (E02H1.2) is an *E. coli* HT115 strain and was a kind gift of Dr. Yelena Budovskaya (SILS, Science Park, University of Amsterdam).

For the fertility assays, hermaphrodite worms were treated with E02H1.2 RNAi from L4 stage of development, and allowed to lay eggs. Their progeny (F1) was continuously exposed to the RNAi treatment, and observed throughout development. The number of F2 progeny from individual F1 worms (N=30) was counted daily for the 4 first days of adulthood. Images of F1 worms before and after reaching adulthood were taken with a Leica DFC320 camera, using an Axio Observer.A1 (Zeiss) microscope and a 10x magnification objective. For imaging, worms were placed on top of a 2% agarose pad on slides, and anaesthetized using 1 mM Levamisole in M9 buffer. For the respiration assays, F1 worms at day 1 of adulthood were used.

## Acknowledgements

The authors thank members of the Houtkooper team as well as Prof. U. Brandt for support and helpful suggestions. Some *C. elegans* strains were provided by the CGC, which is funded by NIH Office of Research Infrastructure Programs (P40 OD010440).

## References

1 Perrault, M., Klotz, B. and Housset, E. (1951) [Two cases of Turner syndrome with deaf-mutism in two sisters]. Bull Mem Soc Med Hop Paris, 67, 79–84.

2 Nishi, Y., Hamamoto, K., Kajiyama, M. and Kawamura, I. (1988) The Perrault syndrome: clinical report and review. Am J Med Genet, 31, 623–629.

3 Gottschalk, M.E., Coker, S.B. and Fox, L.A. (1996) Neurologic anomalies of Perrault syndrome. Am J Med Genet, 65, 274–276.

4 Fiumara, A., Sorge, G., Toscano, A., Parano, E., Pavone, L. and Opitz, J.M. (2004) Perrault syndrome: evidence for progressive nervous system involvement. Am J Med Genet A, 128A, 246–249.

5 Pierce, S.B., Walsh, T., Chisholm, K.M., Lee, M.K., Thornton, A.M., Fiumara, A., Opitz, J.M., Levy-Lahad, E., Klevit, R.E. and King, M.C. (2010) Mutations in the DBP-deficiency protein HSD17B4 cause ovarian dysgenesis, hearing loss, and ataxia of Perrault Syndrome. Am J Hum Genet, 87, 282–288.

6 Pierce, S.B., Chisholm, K.M., Lynch, E.D., Lee, M.K., Walsh, T., Opitz, J.M., Li, W., Klevit, R.E. and King, M.C. (2011) Mutations in mitochondrial histidyl tRNA synthetase HARS2 cause ovarian dysgenesis and sensorineural hearing loss of Perrault syndrome. Proc Natl Acad Sci U S A, 108, 6543–6548.

7 Pierce, S.B., Gersak, K., Michaelson-Cohen, R., Walsh, T., Lee, M.K., Malach, D., Klevit, R.E., King, M.C. and Levy-Lahad, E. (2013) Mutations in LARS2, encoding mitochondrial leucyl-tRNA synthetase, lead to premature ovarian failure and hearing loss in Perrault syndrome. Am J Hum Genet, 92, 614–620.

8 Jenkinson, E.M., Rehman, A.U., Walsh, T., Clayton-Smith, J., Lee, K., Morell, R.J., Drummond, M.C., Khan, S.N., Naeem, M.A., Rauf, B. et al. (2013) Perrault syndrome is caused by recessive mutations in CLPP, encoding a mitochondrial ATP-dependent chambered protease. Am J Hum Genet, 92, 605–613.

9 Ahmed, S., Jelani, M., Alrayes, N., Mohamoud, H.S., Almramhi, M.M., Anshasi, W., Ahmed, N.A., Wang, J., Nasir, J. and Al-Aama, J.Y. (2015) Exome analysis identified a novel missense mutation in the CLPP gene in a consanguineous Saudi family expanding the clinical spectrum of Perrault Syndrome type-3. J Neurol Sci, 353, 149–154.

10 Morino, H., Pierce, S.B., Matsuda, Y., Walsh, T., Ohsawa, R., Newby, M., Hiraki-Kamon, K., Kuramochi, M., Lee, M.K., Klevit, R.E. et al. (2014) Mutations in Twinkle primase-helicase cause Perrault syndrome with neurologic features. Neurology, 83, 2054–2061.

11 Uchiumi, T., Ohgaki, K., Yagi, M., Aoki, Y., Sakai, A., Matsumoto, S. and Kang, D. (2010) ERAL1 is associated with mitochondrial ribosome and elimination of ERAL1 leads to mitochondrial dysfunction and growth retardation. Nucleic Acids Res, 38, 5554–5568.

12 Dennerlein, S., Rozanska, A., Wydro, M., Chrzanowska-Lightowlers, Z.M. and Lightowlers, R.N. (2010) Human ERAL1 is a mitochondrial RNA chaperone involved in the assembly of the 28S small mitochondrial ribosomal subunit. Biochem J, 430, 551–558.

13 Tu, C., Zhou, X., Tropea, J.E., Austin, B.P., Waugh, D.S., Court, D.L. and Ji, X. (2009) Structure of ERA in complex with the 3' end of 16S rRNA: implications for ribosome biogenesis. Proc Natl Acad Sci U S A, 106, 14843–14848.

14 Heide, H., Bleier, L., Steger, M., Ackermann, J., Drose, S., Schwamb, B., Zornig, M., Reichert, A.S., Koch, I., Wittig, I. et al. (2012) Complexome profiling identifies TMEM126B as a component of the mitochondrial complex I assembly complex. Cell Metab, 16, 538–549.

15 Houtkooper, R.H., Mouchiroud, L., Ryu, D., Moullan, N., Katsyuba, E., Knott, G., Williams, R.W. and Auwerx, J. (2013) Mitonuclear protein imbalance as a conserved longevity mechanism. Nature, 497, 451–457.

16 Sijen, T., Fleenor, J., Simmer, F., Thijssen, K.L., Parrish, S., Timmons, L., Plasterk, R.H. and Fire, A. (2001) On the role of RNA amplification in dsRNA-triggered gene silencing. Cell, 107, 465–476.

17 Szczepanowska, K., Maiti, P., Kukat, A., Hofsetz, E., Nolte, H., Senft, K., Becker, C., Ruzzenente, B., Hornig-Do, H.T., Wibom, R. et al. (2016) CLPP coordinates mitoribosomal assembly through the regulation of ERAL1 levels. EMBO J, 35, 2566–2583.

18 Crimi, M., Galbiati, S., Perini, M.P., Bordoni, A., Malferrari, G., Sciacco, M., Biunno, I., Strazzer, S., Moggio, M., Bresolin, N. et al. (2003) A mitochondrial tRNA(His) gene mutation causing pigmentary retinopathy and neurosensorial deafness. Neurology, 60, 1200–1203.

19 Kokotas, H., Petersen, M.B. and Willems, P.J. (2007) Mitochondrial deafness. Clin Genet, 71, 379–391.

20 Guan, M.X. (2004) Molecular pathogenetic mechanism of maternally inherited deafness. Ann N Y Acad Sci, 1011, 259–271.

21 Thouas, G.A., Trounson, A.O. and Jones, G.M. (2006) Developmental effects of sublethal mitochondrial injury in mouse oocytes. Biol Reprod, 74, 969–977.

22 Igosheva, N., Abramov, A.Y., Poston, L., Eckert, J.J., Fleming, T.P., Duchen, M.R. and McConnell, J. (2010) Maternal diet-induced obesity alters mitochondrial activity and redox status in mouse oocytes and zygotes. PLoS One, 5, e10074.

23 Ben-Meir, A., Yahalomi, S., Moshe, B., Shufaro, Y., Reubinoff, B. and Saada, A. (2015) Coenzyme Q-dependent mitochondrial respiratory chain activity in granulosa cells is reduced with aging. Fertil Steril, 104, 724–727.

24 Dillin, A., Hsu, A.L., Arantes-Oliveira, N., Lehrer-Graiwer, J., Hsin, H., Fraser, A.G., Kamath, R.S., Ahringer, J. and Kenyon, C. (2002) Rates of behavior and aging specified by mitochondrial function during development. Science, 298, 2398–2401.

25 Lee, S.S., Lee, R.Y., Fraser, A.G., Kamath, R.S., Ahringer, J. and Ruvkun, G. (2003) A systematic RNAi screen identifies a critical role for mitochondria in C. elegans longevity. Nat Genet, 33, 40–48.

26 Rea, S.L., Ventura, N. and Johnson, T.E. (2007) Relationship between mitochondrial electron transport chain dysfunction, development, and life extension in Caenorhabditis elegans. Plos Biology, 5, 2312–2329.

27 Arnold, K., Bordoli, L., Kopp, J. and Schwede, T. (2006) The SWISS-MODEL workspace: a web-based environment for protein structure homology modelling. Bioinformatics, 22, 195–201.

28 Janssen, A.J., Trijbels, F.J., Sengers, R.C., Smeitink, J.A., van den Heuvel, L.P., Wintjes, L.T., Stoltenborg-Hogenkamp, B.J. and Rodenburg, R.J. (2007) Spectrophotometric assay for complex I of the respiratory chain in tissue samples and cultured fibroblasts. Clin Chem, 53, 729–734.

29 Wittig, I., Braun, H.P. and Schagger, H. (2006) Blue native PAGE. Nat Protoc, 1,418428.

30 Huynen, M.A., Muhlmeister, M., Gotthardt, K., Guerrero-Castillo, S. and Brandt, U. (2015) Evolution and structural organization of the mitochondrial contact site (MICOS) complex and the mitochondrial intermembrane space bridging (MIB) complex. Biochim Biophys Acta, 1863, 91–101.

31 Cox, J. and Mann, M. (2008) MaxQuant enables high peptide identification rates, individualized p.p.b.-range mass accuracies and proteome-wide protein quantification. Nat Biotechnol, 26, 1367–1372.

32 Giese, H., Ackermann, J., Heide, H., Bleier, L., Drose, S., Wittig, I., Brandt, U. and Koch, I. (2015) NOVA: a software to analyze complexome profiling data. Bioinformatics, 31, 440–441.

33 Brenner, S. (1974) The genetics of Caenorhabditis elegans. Genetics, 77, 71–94.

34 Kamath, R.S., Martinez-Campos, M., Zipperlen, P., Fraser, A.G. and Ahringer, J. (2001) Effectiveness of specific RNA-mediated interference through ingested doublestranded RNA in Caenorhabditis elegans. Genome Biol, 2, RESEARCH0002.

